# Gene Expression in Lung Epithelial Cells Following Interaction with *Pneumocystis carinii* and its Specific Life Forms Yields Insights into Host Gene Responses to Infection

**DOI:** 10.1101/2021.11.22.469523

**Authors:** Theodore J. Kottom, Eva M. Carmona, Andrew H. Limper

**Affiliations:** Thoracic Diseases Research Unit, Departments of Medicine and Biochemistry, Mayo Clinic College of Medicine, Rochester, Minnesota, 55905. USA

**Keywords:** A549, epithelium, microarray, *Pneumocystis*

## Abstract

*Pneumocystis* spp. interacts with epithelial cells in the alveolar spaces of the lung. It is thought that the binding of *Pneumocystis* to host cell epithelium is needed for life cycle completion and proliferation. The effect of this interaction on lung epithelial cells have previously shown that the trophic form of this organism greatly inhibits p34^*cdc2*^ activity, a serine/threonine kinase required for transition from G_2_ to M phase in the cell cycle. To gain further insight into the host response during Pneumocystis pneumonia (PCP), we used microarray technology to profile epithelial cell (A549) gene expression patterns following *Pneumocystis carinii* interaction. Furthermore, we isolated separate populations of cyst and trophic forms of *P. carinii*, which were then applied to the lung epithelial cells. Differential expression of genes involved in various cellular functions dependent on the specific *P. carinii* life form in contact with the A549 cell were identified. The reliability of our data was further confirmed by Northern blot analysis on a number of selected up or down regulated transcripts. The transcriptional response to *P. carinii* was dominated by cytokines, apoptotic, and anti-apoptotic related genes. These results reveal several previously unknown effects of *P. carinii* on the lung epithelial cell and provide insight into the complex interactions of host and pathogen.

## INTRODUCTION

Pneumocystis jirovecii pneumonia (PJP) is a fungal infection caused by *Pneumocystis jirovecii* and affects those who are immunocompromised, especially those with AIDS [1, 2]. Immune and cytokine response to *Pneumocystis* has been well studied in the host and macrophage [2-9]. However, little is known about the responses generated by this organism on the epithelial cell level, especially at the transcriptional level. In the respiratory airway, epithelial cells are active participants in the early inflammatory response to defend the airway [10]. Due to their location in the lung, epithelial cells are often the first cells to encounter potential microbial pathogens.

Previous evidence suggests that *Pneumocystis* adherence to lung epithelial cells maybe a stimulus for the organism to replicate [11]. Indeed, the *Pneumocystis* trophic life form when subjected in vitro to extracellular matrixes present on host cells such as collagen or fibronectin causes an increase in *PCSTE20* transcription, a molecule shown to be important in yeast mating [11]. This interaction of *Pneumocystis* with host epithelium has caused the induction of numerous cytokines including intercellular adhesin molecule-1 (ICAM-1), interleukin-8 (IL-8), macrophage inflammatory protein-2 (MIP-2), and monocyte chemoattractant protein 1 (MCP-1) [12-14]. Although this data demonstrated that *Pneumocystis* indeed cause release of these cytokines, the overall molecular mechanisms that regulate their expression remains unclear. Furthermore, an in-depth profile of host genes whose expression is modulated by *Pneumocystis* including signal transduction pathways causing these changes remain largely unknown.

A number of studies have examined host/pathogen interaction by measuring host transcriptional profiles using high-density microarrays [15-18]. These studies have enhanced our knowledge of host responses to various pathogens of eukaryotic, prokaryotic, as well as viral in nature [16-25]. The goal of our current study was to examine the global changes in transcriptional responses of lung epithelial cells to early infection with *Pneumocystis* mixed as well as isolated populations of cyst and trophic forms. We used the A549 lung epithelial cell line as a model system since this line strongly supports *Pneumocystis* binding of the organisms and has been used extensively and is the most cited epithelial cell line used for Pneumocystis interactions [26-34]. Interestingly, by isolating separate populations of cyst versus trophic forms, we were able to dissect out unique transcriptional responses indicating that components and/or interactions of these life forms can cause different host responses. To confirm our microarray results, the expression profile of some of the differently regulated genes were examined by Northern blot and showed a close correlation with the microarray results. Lastly, we showed that Bfl-1/A1, a member of the Bcl-2 family of proteins involved in regulating the apoptotic response [35] is activated in lung epithelial cells upon *P. carinii* contact and that this activation is NF-κB dependent.

## METHODS

### Cell lines and strains

The human lung epithelium A549 cell line (CCL-185™) was obtained from ATCC®. The cell lines were grown in Ham’s F12K medium with 2 mM L-glutamine adjusted to contain 1.5 g/L sodium bicarbonate, supplemented with 10% fetal bovine serum.

### Isolation of *Pneumocystis carinii*

All animal experiments were conducted in accordance with the guidelines of the Mayo Institutional Animal Care and Use Committee (IACUC). Pneumocystis pneumonia (PCP) was induced in rats immunosuppressed as previously described [36]. *Pneumocystis carinii* organisms were derived from the American Type Culture Collection (ATCC^®^). Briefly, whole populations of *P. carinii* containing both trophic forms and cysts were purified from chronically infected rat lungs by homogenization and filtration through 10-µm filters, and further fractionated into enriched populations of trophic forms (99.5% pure) and cysts (>40-fold enriched) by differential filtration through 3-µm filters, as previously described [37]. If other microorganisms were noted in the lavage smear or microbiological culture, the material was discarded. Previously, this method of organism isolation yielded <0.125U/ml of soluble endotoxin using a sensitive Limulus amoebocyte lysate assay as previously described [38].

### Cell infection and total RNA preparation

Approximately 1 × 10^6^ A549 cells were infected with *P. carinii* organisms at a 75:1 multiplicity of infection for 3.0 h. *P. carinii* organisms were applied either directly on cells directly or on Transwell inserts above the cells to distinguish between transcription profiles created by soluble factors released from *P. carinii* versus profiles created by actual organism-to-cell contact. Upon application of *P. carinii* organisms to either cells or Transwell inserts, plates were spun at 500 x g 5.0 minutes to synchronize infection. After 3.0 hours, total RNA was extracted with the RNeasy® Mini Kit (Qiagen, Valencia, CA).

### Hybridization to oligonucleotide chips

Oligonucleotide GeneChip™ Human Genome U133A 2.0 Array (ThermoFisher Scientific, Waltham MA) are oligonucleotide microarrays, which allow the characterization of genome-wide expression levels of approximately 18,400 transcripts and variants, including 14,500 well-characterized human genes. Each 25-mer oligonucleotide probe is uniquely complementary to a gene with approximately 16 pairs of oligonucleotide probes used to measure the transcript level of each of the sequences represented. The Mayo Clinic Cancer Center Microarray Facility conducted labeling and hybridization of probes to microarrays. For our experiments, we were interested and report genes that had a 4-fold change from untreated control samples.

### Northern blots

Total RNA (5 μg) was separated by agarose gel electrophoresis using standard molecular biological techniques. Northern blots were conducted following standard procedures using PCR-generated probes created from primers listed in Table I. Probes were labeled with α- ^32^P dATP (Sigma Aldrich, St. Louis, MO). Following hybridization with ExpressHyb solution (Takara) membranes were washed 4 times and visualized by autoradiography. Blots were also subjected to stripping and re-probing according to ExpressHyb protocol.

**TABLE I.** Clustering of differently expressed genes with *P. carinii* on Transwell inserts. Expression profile of epithelial cell gene expression with *P. carinii* in Transwell inserts above A549 cells but not in direct contact with the lung cells were conducted in two independent experiments and average fold change (FC) reported.

| Accession number | Fold Change (FC) | Gene |
| --- | --- | --- |
| AI608725 | 3.3 | Similar to intercellular adhesion molecule 1 (*ICAM1*) |
| M37435 | 5.3 | Colony stimulating factor (*CSF-1*) |
| NM_002198 | 5.6 | Interferon regulatory factor (*IRF1*) |
| M15329 | 5.9 | Interleukin 1-α (*IL1A*) |
| NM_015714 | 6.4 | G<sub>0</sub>/G<sub>1</sub> switch 2 (*GOS2*) |
| NM_000963 | 6.9 | Prostaglandin-endoperoxide synthase 2 (*COX2*) |
| NM_025079 | 7.2 | Unknown protein |
| NM_006291 | 7.3 | TNFα-induced protein 2 (*TNFAIP2*) |
| NM_000584 | 7.9 | Interleukin 8 (*IL-8*) |
| BC002666 | 8.4 | Guanylate binding protein 1 |
| NM_005204 | 8.4 | *MAP3K8* |
| NM_004428 | 9.1 | Ephrin-A1 (*EFNA1*) |
| AI078167 | 9.4 | MHC enhancer binding protein |
| NM_006290 | 12.1 | TNFα-induced protein 3 (*TNFAIP3*) |
| AI738896 | 12.3 | Similar to human DNA-binding protein A20 |
| NM_001511 | 13 | *CXCL1* |
| U37546 | 14.2 | IAP homolog C (*MIHC*) |
| AF043337 | 16.3 | *IL8* variant |
| NM_002852 | 16.7 | Pentraxin-related gene (*PTX3*) |
| M57731 | 26 | *CXCL2* |
| S69738 | 31.5 | *MCP-1* |
| NM_002090 | 44.1 | *GRO3* |
| NM_003897 | 59.2 | *GM-CSF* |
| NM_004591 | 167 | *CCL20* |

### Analysis of Bfl-1 promoter activity

A549 cells (5 ×10^5^) were incubated overnight with wildtype plasmid *Bfl-1*-CAT [35] in Lipofectamine™ 2000 in OPTI-MEM® reduced-serum media (ThermoFisher Scientific). Empty pCMV (ThermoFisher Scientific) was used a negative empty control vector. The following day, media was removed, and 2.0 ml fresh media was added back. The *P. carinii* infection scheme listed above, including polymyxin B-treated organisms was employed and the plates were placed at 37^0^C/5%C0_2_ for 6 hours. At 6 hours, cell lysates were harvested, and CAT assays performed according to a CAT enzyme assay system (Promega, Fitchburg, WI).

### Statistical analysis

For multigroup data, initial analysis was first performed with analysis of variance (ANOVA) to determine overall different differences. If ANOVA indicated overall differences, subsequent group analysis was then performed by 2-sample Student’s *t*-test for normally distributed variables. Evaluation of data was conducted using Prism 9 for macOS, version 9.1.2 (GraphPad, San Diego, CA). Values of *p* < 0.05 were considered significant.

## RESULTS

### Study of gene expression profiles by microarray after binding of the A549 cell line with *Pneumocystis carinii* mixed populations and isolated trophic forms and cysts

We used GeneChip™ Human Genome U133A 2.0 Arrays to monitor mRNA levels in a lung epithelial cell line (A549) after challenge with *Pneumocystis carinii* for 3 hours. Transcript abundance from approximately 18,400 human genes was analyzed in duplicate experiments. Non-infected A549 cells were used as a control. For the purpose of this paper, we report only those genes that exhibited greater than a 4-fold change in steady-state mRNA abundance after 3 hours of infection. Infection of the epithelial cell line induced marked change in host cell gene expression. For the 4 treatment groups, total number of genes with 4-fold changes were as follows: *P. carinii* in Transwells above A549 cells (*P. carinii* not in contact with A549); 24 genes, isolated *P. carinii* cysts forms in contact with A549 cells; 26 genes, isolated *P. carinii* trophic forms in contact with A549; 48 genes, and finally, mixed populations of *P. carinii* cyst and trophic forms in contacts with A549 lung cells; 70 genes, respectively. This distribution of the two independent experiments is illustrated in (Figure 1).

**Figure 1.**
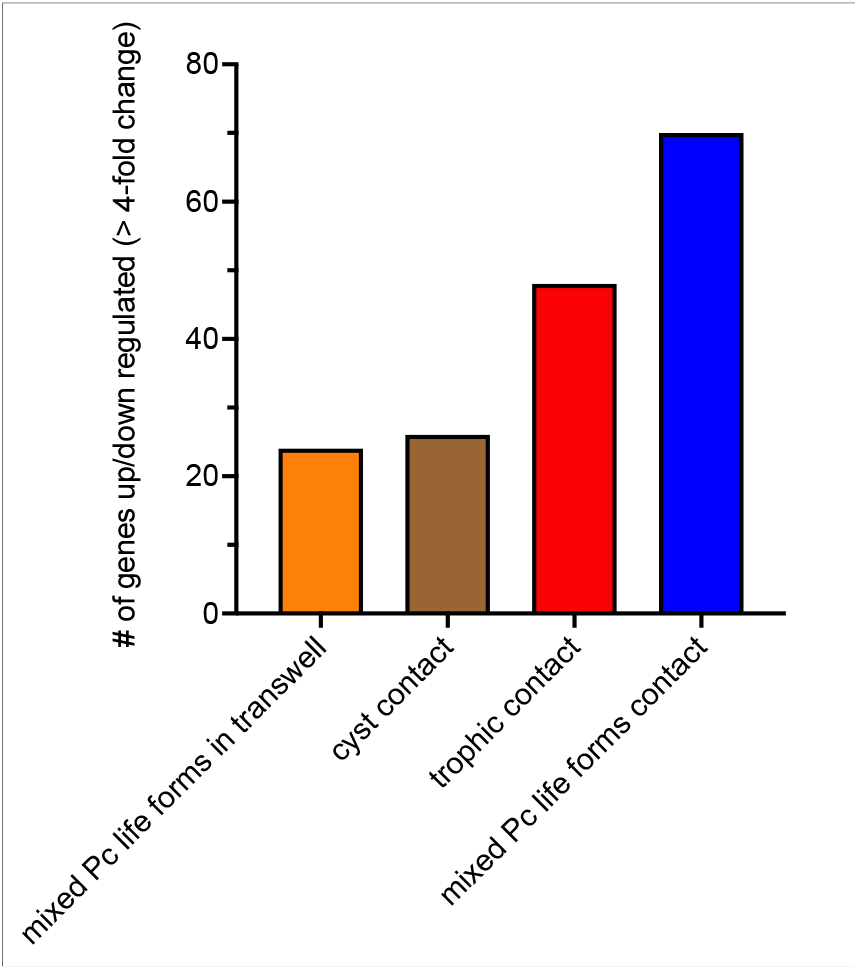
Bar graph of equal to or 4-fold up-or downregulated transcripts in A549 epithelial cells (from two independent experiments) following the listed *P. carinii* (Pc) organism treatment groups versus mock infection with the Oligonucleotide GeneChip™ Human Genome U133A 2.0 Array. A total list of each of these genes for each of the treatment groups is listed in Table I.-Table V.

To further confirm the microarray findings, we conducted Northern blot analysis of identified transcripts. Indeed, upon analyzing eight randomly differentially expressed representative genes, we found a close correlation between the fold change data from the microarray experiments and the intensities in the Northern blot measurements. Figure 2 illustrates this comparison.

**Figure 2.**
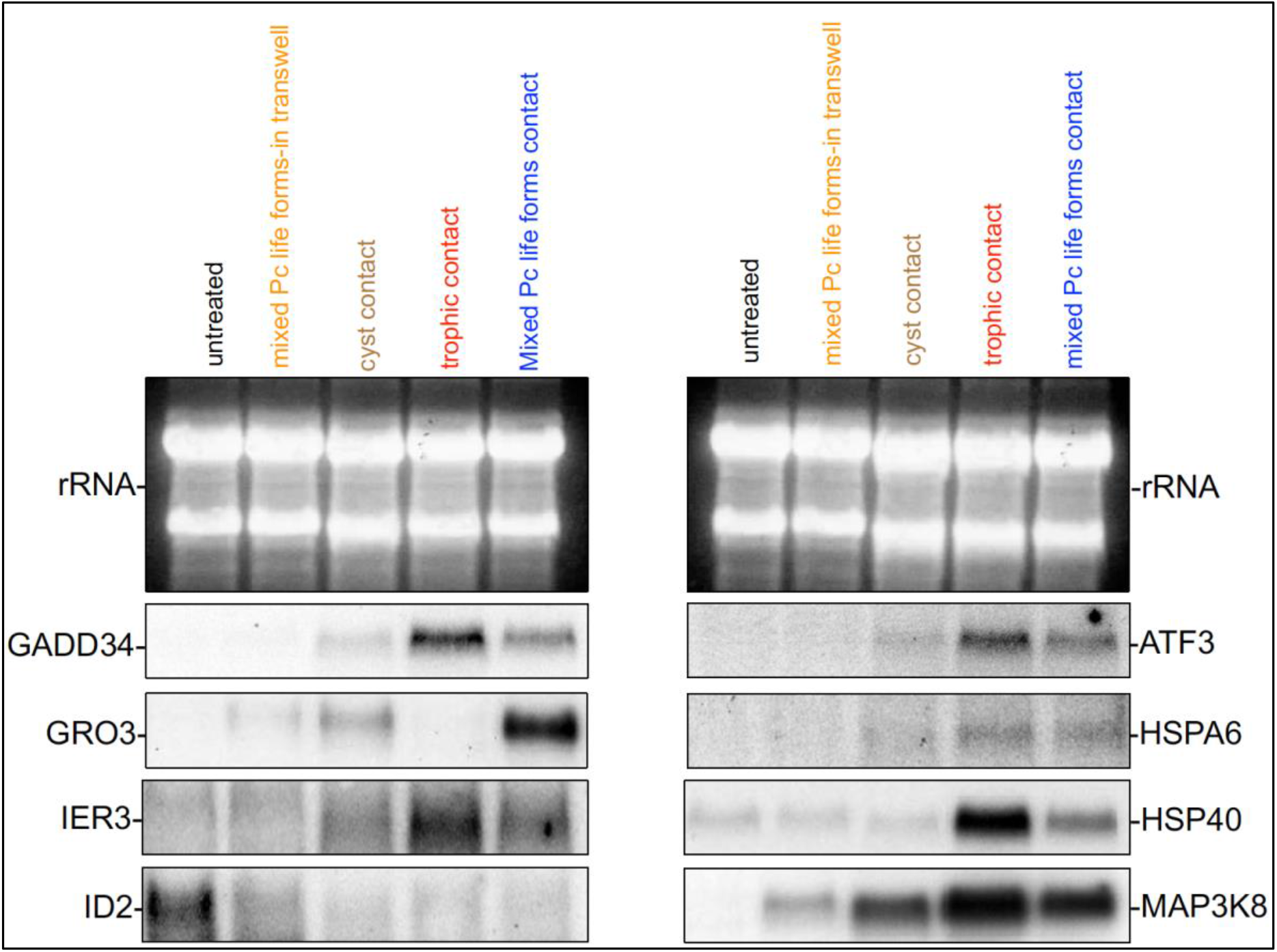
Comparison of Northern blotting results for eight representative differentially expressed genes determined in the microarray survey. Northern hybridization of mRNA from the respective *P. carinii* (Pc) and epithelial interactions. PCR-generated probes to specific gene targets listed were generated with the primers listed in Supplementary Table I. The top photograph in each panel demonstrates the two A459 cell major ribosomal subunits demonstrating equal RNA loading.

Further validation of the results was provided by finding gene expression changes that were previously known to be modulated upon *P. carinii* interactions with A549 cells. Among the various genes listed in tables II-V. found to be up-regulated at least 4-fold over the control, at least 3 genes have been previously described as *Pneumocystis* induced. For example, ICAM-1, IL-8, and MCP-1 protein production have already been reported to be induced following *Pneumocystis* infection of the A549 cell line [13, 14, 39]. The other genes that we found in our current study are, to the best of our knowledge, newly described transcriptional responses to *Pneumocystis* interactions with the A549 lung epithelial cell line.

**TABLE II.** Clustering of differently expressed genes with *P. carinii* cyst forms in contact with A549 cell line. Expression profile of epithelial cell gene expression with *P. carinii* cyst forms on A549 cells were conducted in two independent experiments and average fold change (FC) reported.

| Accession number | Fold Change (FC) | Gene |
| --- | --- | --- |
| NM_002166 | -6.0 | DNA inhibitor of binding 2 (*ID2*) |
| AL021977 | 4.4 | Unknown protein |
| W46388 | 4.9 | Superoxide dismutase precursor 2 (*SOD2*) |
| NM_007203 | 5.1 | PALM2 and AKAP2 fusion<br>(*PALM2AKAP2*) |
| NM_025079 | 5.3 | Unknown protein |
| NM_000963 | 6.0 | Prostaglandin-endoperoxide synthase 2<br>(*COX2*) |
| NM_000201 | 6.1 | Intercellular adhesion<br>molecule 1 (*ICAM1*) |
| AI078167 | 6.2 | MHC enhancer binding<br>protein |
| NM_006291 | 6.5 | TNF-α induced protein 2<br>(*TNFAIP2*) |
| NM_005204 | 7.6 | *MAP3K8* |
| AF078844 | 8 | Unknown protein |
| NM_004428 | 8.2 | Ephrin-A1 (*EFNA1*) |
| BF246115 | 8.9 | Unknown protein |
| NM_000584 | 9.4 | Interleukin 8 (*IL8*) |
| NM_006290 | 9.8 | TNFα-induced protein 3<br>(*TNFAIP3*) |
| NM_001511 | 11.2 | *GRO1* |
| AI738896 | 11.5 | Similar to human DNA-binding protein A20 |
| U37546 | 15.3 | IAP homolog C (*MIHC*) |
| NM_002852 | 16.7 | Pentraxin-related gene<br>(*PTX3*) |
| AF043337 | 17.1 | *IL-8* variant |
| S69738 | 19.2 | *MCP-1* |
| M57731 | 20.2 | *CXCL2* |
| NM_002090 | 28.9 | *GRO3* |
| M11734 | 46.8 | *GM-CSF* |
| NM_004049 | 47.5 | BCL2-related protein A1<br>(*BFL-1/A1*) |
| NM_004591 | 92 | *CCL20* |

### Identification of specific genes modulated by isolated *P. carinii* life forms or mixed populations

By using the differential filtration method, we reported previously to isolate specific life forms of *P. carinii* [37], we were able obtain transcriptional responses for each life form directly interacting with A549 cells. A number of these gene responses were noted to be upregulated upon contact with all of the life forms presented to the cell line or *P. carinii* in Transwells such as *EFNA1, GRO1*, and *CCL20*. Although a number of up-regulated genes were noted under all conditions consisting of mixed *P. carinii* populations or both separated forms, we were able to determine specific transcripts either up or down-regulated based on contact with a specific *P. carinii* life form. Transcripts of interest in response to the cyst form (Table V) include *ID2* a helix-loop-helix (HLH) transcription regulator thought to be important in cell growth and differentiation [40] which was down-regulated in *P. carinii* cyst binding to A549 cells (Figure 2 and Table V.), as well as human superoxide dismutase precursor 2 (*SOD2*) (Accession W46388). Finally, unique to cyst form exposure were the up regulation of two unknown genes in the A549 cells, namely AF078844 and BF246115 (Table V.) with nucleotide similarity to a family of cysteine-rich, metal binding intracellular proteins, which have been linked to cell proliferation [41].

**TABLE III.** Clustering of differently expressed genes with *P. carinii* trophic forms in contact with A549 cells. Expression profile of epithelial cell gene expression with *P. carinii* trophic forms on A549 cells were conducted in two independent experiments and average fold change (FC) reported.

| Accession number | Fold Change (FC) | Gene |
| --- | --- | --- |
| NM_002090 | -5.3 | *GRO3* |
| NM_030952 | 4.5 | NUAK family kinase<br>(*NUAK2*) |
| NM_025079 | 4.7 | Unknown protein |
| BC004490 | 4.8 | v-fos FBJ viral oncogene<br>homolog |
| S69738 | 5.3 | *MCP-1* |
| BG537255 | 5.6 | Unknown protein |
| AF043337 | 5.6 | *IL-8* variant |
| X51757 | 5.8 | Heat-shock protein 70<br>(HSP70) |
| AI935096 | 6.3 | Unknown protein |
| NM_001511 | 6.4 | GRO1 |
| NM_004281 | 7.7 | BAG cochaperone 3(BAG3) |
| NM_002135 | 8 | Nuclear receptor subfamily<br>4 group A member 1<br>(NR4A1) |
| NM_006186 | 8.4 | Nuclear receptor subfamily<br>4 group A member 2<br>(NR4A2) |
| AF133207 | 8.7 | Protein kinase (H11) |
| NM_006145 | 9.2 | Heat-shock protein 40<br>(HSP40) |
| M57731 | 9.3 | CXCL2 |
| AA530892 | 9.4 | MAP kinase phosphatase-1<br>(MKP-1) |
| U37546 | 9.7 | IAP homolog C (MIHC) |
| NM_002852 | 10 | Pentraxin-related gene<br>(PTX3) |
| NM_003407 | 10.7 | ZFP36 ring finger protein<br>(ZFP36) |
| AI078167 | 11.5 | MHC enhancer-binding<br>protein |
| U69563 | 12.1 | Unknown protein |
| NM_000963 | 13.2 | Prostaglandin-endoperoxide synthase 2<br>(COX2) |
| NM_001674 | 13.6 | Activating transcription<br>factor 3 (ATF3) |
| NM_002228 | 13.7 | v-jun avian sarcoma virus<br>homolog |
| BE675435 | 13.9 | DNA-binding protein |
| M11734 | 14.9 | GM-CSF |
| NM_003897 | 15.7 | Early response 3(IER3) |
| AL021977 | 17.4 | Unknown protein |
| NM_001554 | 17.7 | Cellular communication<br>network factor 1 (CCN1) |
| NM_006290 | 17.7 | TNFα-induced protein 3<br>(TNFAIP3) |
| NM_014330 | 17.8 | Protein phosphatase 1<br>regulatory subunit 15A<br>(PPP1R15A) |
| NM_000584 | 19.6 | IL-8 |
| NM_005204 | 21.0 | MAP3K8 |
| NM_004428 | 21.1 | Ephrin-A1 (EFNA1) |
| NM_004049 | 25.4 | BCL2-related protein A1<br>(BFL-1/A1) |
| AI738896 | 25.8 | Similar to human DNA-binding protein A20 |
| AB017493 | 26.2 | DNA-binding zinc finger (*GBF*) |
| AF003114 | 28.7 | *CYR61* |
| NM_005346 | 32.9 | Heat-shock protein 1B (*HSP1B*) |
| D13889 | 35.4 | *Id-1H* |
| S77154 | 39.7 | NGFI-B transcription factor (*TINUR*) |
| U83981 | 52 | Growth arrest and DNA damage 34 (*GADD34*) |
| U16996 | 55.6 | Protein tyrosine phosphatase |
| NM_004591 | 84.5 | Chemokine ligand 20 (*CCL20*) |
| M15329 | 143.2 | Interleukin 1-α (*IL1A*) |
| AL556438 | 412 | Unknown protein |
| NM_005345 | 442 | Heat-shock protein 1A (*HSP1A*) |

**TABLE IV.** Clustering of differently expressed genes with *P. carinii* mixed life forms in contact with A549 cell line. Expression profile of epithelial cell gene expression with *P. carinii* total life forms on A549 cells were conducted in two independent experiments and average fold change (FC) reported.

| Accession number | Fold Change (FC) | Gene |
| --- | --- | --- |
| AL050388 | 4.2 | Manganese superoxide dismutase |
| AF078077 | 4.5 | Growth arrest and DNA damage 45 beta (*GADD45β*) |
| BG054844 | 4.8 | Unknown protein |
| AI572079 | 5.1 | Unknown protein |
| NM_030952 | 5.4 | NUAK family kinase (*NUAK2*) |
| U16996 | 5.5 | Protein tyrosine phosphatase |
| NM_005195 | 5.7 | CCAAT enhancer binding protein (*CEBPD*) |
| NM_015714 | 6.1 | lymphocyte G<sub>0</sub>/G<sub>1</sub> (*G0S2*) |
| NM_003364 | 6.6 | Uridine phosphorylase (*UPP1*) |
| NM_002155 | 6.6 | Heat shock protein 6 (*HSPA6*) |
| NM_005931 | 6.7 | MHC class I polypeptide (*MICB*) |
| M24779 | 6.7 | Protein kinase (*PIM1*) |
| BG537255 | 6.9 | Unknown protein |
| NM_004281 | 7 | BCL2 (*BAG3*) |
| NM_0015675 | 7.1 | Unknown protein |
| NM_006145 | 7.5 | Heat-shock protein<br>(*HSP40*) |
| BG491844 | 7.5 | Unknown protein |
| NM_007203 | 7.9 | A kinase (*PRKA*) anchor<br>protein 2 |
| AF133207 | 7.9 | Protein kinase (*H11*) |
| NM_006186 | 8.2 | Nuclear receptor subfamily<br>4 group A member 2<br>(*NR4A2*) |
| NM_005261 | 8.7 | GTP-binding protein (*GEM*) |
| NM_025079 | 8.8 | Unknown protein |
| NM_005346 | 9 | Heat-shock protein 1B<br>(*HSP1B*) |
| NM_001924 | 9 | Growth arrest and DNA<br>damage 45 beta<br>(*GADD45α*) |
| NM_000963 | 9.3 | Prostaglandin-endoperoxide synthase 2<br>(*COX2*) |
| NM_003897 | 9.4 | Early response 3(*IER3*) |
| NM_005345 | 9.8 | Heat-shock protein 1A<br>(*HSP1A*) |
| AI935096 | 10.3 | Unknown protein |
| NM_000584 | 10.4 | *IL-8* |
| AF003114 | 10.5 | Carnitine O-octanoyltransferase<br>(*CYR61*) |
| NM_001945 | 11 | Heparin binding EGF like<br>growth factor (*HBEGF*) |
| S77154 | 11.1 | NGFI-B transcription factor<br>(*TINUR*) |
| NM_001554 | 11.3 | Cellular communication<br>network factor 1 (*CCN1*) |
| U69563 | 11.5 | Unknown protein |
| NM_002198 | 12.8 | Interferon regulatory factor<br>(*IRF1*) |
| NM_001993 | 14.8 | Coagulation factor III, tissue<br>factor (F3), transcript<br>variant 1 |
| M15329 | 14.9 | Interleukin 1-alpha (*IL1A*) |
| NM_003407 | 16.1 | ZFP36 ring finger protein<br>(*ZFP36*) |
| NM_002341 | 16.3 | Lymphotoxin $\beta$ (*LTβ*) |
| AF043337 | 16.3 | *IL-8* variant |
| NM_005204 | 16.4 | *MAP3K8* |
| NM_002228 | 16.5 | v-jun sarcoma oncogene homolog |
| AB017493 | 16.6 | DNA-binding zinc finger (*GBF*) |
| NM_001511 | 17.4 | *GRO1* |
| AI078167 | 17.7 | MHC enhancer-binding protein |
| U37546 | 19.4 | IAP homolog C (*MIHC*) |
| NM_001956 | 20.7 | Endothelin 2 (*EDN2*) |
| NM_014330 | 21.2 | Protein phosphatase 1 regulatory subunit 15A (*PPP1R15A*) |
| NM_006732 | 21.6 | Osteosarcoma viral oncogene( *FOSB*) |
| NM_004428 | 22 | Ephrin-A1 (*EFNA1*) |
| AJ224869 | 22.8 | *CXCR4* |
| D49728 | 23.7 | *NAK1* |
| NM_0006001 | 26.1 | *IL-6* |
| NM_001674 | 26.2 | Activating transcription factor 3 (*ATF3*) |
| U83981 | 26.2 | Growth arrest and DNA damage 34 (*GADD34*) |
| M21121 | 27.4 | T cell specific (*RANTES*) |
| X51757 | 30.5 | Heat-shock protein 70 (*HSP70*) |
| NM_006290 | 31.5 | TNFα-induced protein 3 (*TNFAIP3*) |
| AI738896 | 33.3 | Similar to human DNA-binding protein A20 |
| BC002646 | 38.1 | v-jun sarcoma oncogene homolog |
| BC004490 | 40.2 | v-fos FBJ viral oncogene homolog |
| NM_002852 | 42.2 | Pentraxin-related gene (*PTX3*) |
| NM_002135 | 42.4 | Nuclear receptor subfamily 4 group A member 1 (*NR4A1*) |
| S69738 | 48.7 | *MCP-1* |
| BE675435 | 59.1 | DNA-binding protein |
| M57731 | 60.2 | *CXCL2* |
| NM_002090 | 84.8 | *GRO3* |
| NM_004049 | 101.6 | BCL2-related protein A1 (*Bfl-1/A1*) |
| M11734 | 227 | *GM-CSF* |
| NM_004591 | 710 | Chemokine ligand 20 (*CCl20*) |

**TABLE V.**
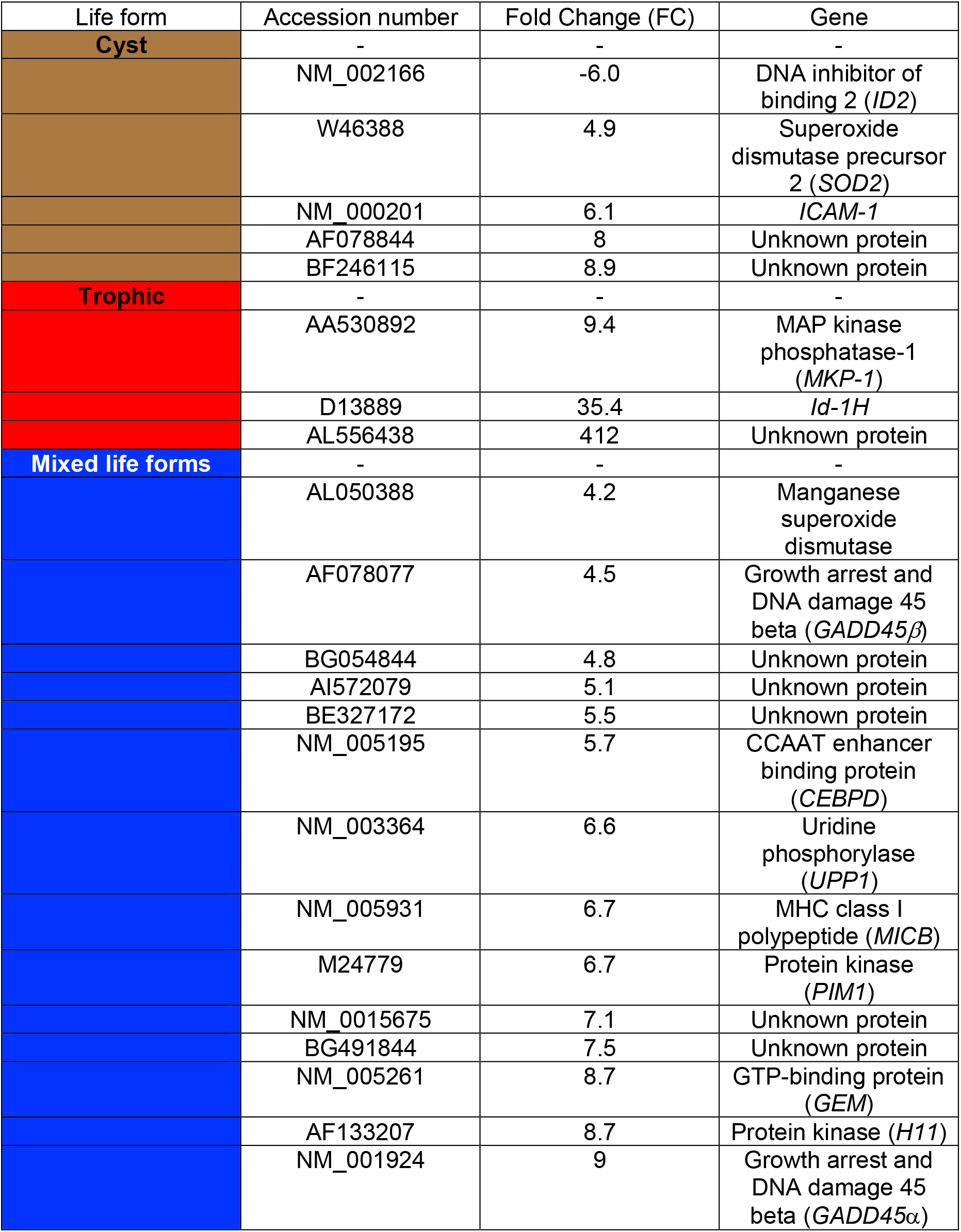
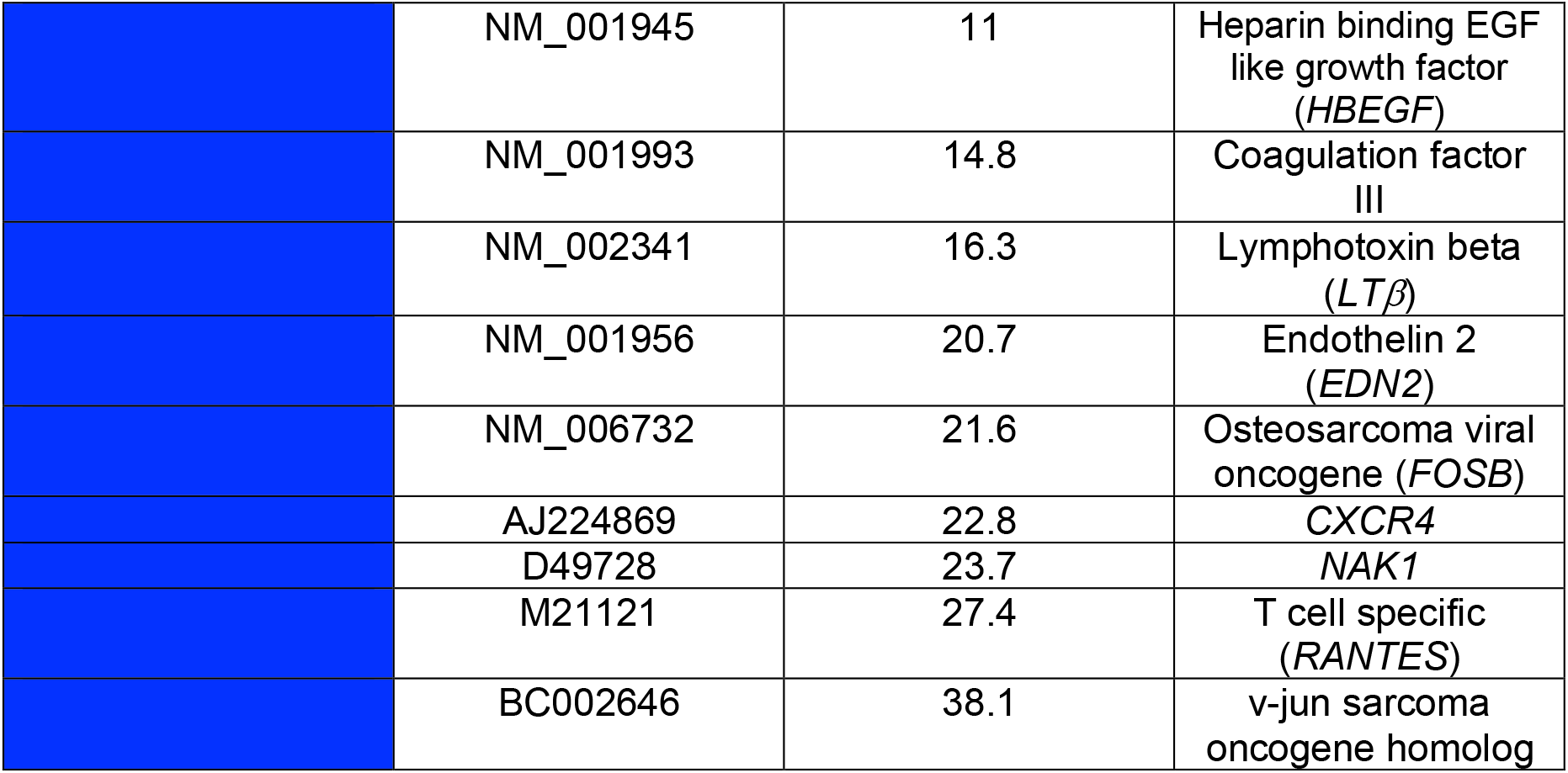
Clustering of differently expressed genes dependent on specific-life forms directly interacting with A549 cells. Expression profile of epithelial cell gene expression with *P. carinii* life forms on A549 cells were conducted in two independent experiments and average fold change (FC) reported.

Next, in the isolated trophic forms in contact with A549 cells, 3 genes were discovered to be modulated upon specific trophic form contact (Table V.). These include one unknown gene/protein product, AL556438, as well as another HLH transcript, *ID-1H* which contrary to the HLH regulator *ID2* which was down-regulated following cyst contact with A549 epithelial cells, was found to be induced in A549 cells upon trophic contact. Most importantly though was the induction of Mitogen-activated protein kinase (MAPK) phophatase-1 (*MKP-1*). MKP-1 is a member of the family of dual specificity protein phosphatases than can dephosphorylate phosphothreonine and phosphotyrosine residues, causing inactivation of MAPKs [42]. It has been shown MKP-1 is a target of the tumor suppressor p53 protein, which suppresses cell growth by inducing cell cycle arrest or apoptosis [43]. Overexpression of MKP-1 prevents cells from entering into the cell cycle [43].

Lastly, we identified a number of genes which provide a unique epithelial cell synergistic response to a mixed population of both *P. carinii* cyst as well as trophic forms (Table V.). Some of these transcripts include *RANTES*, a member of the C-C chemokine subfamily involved in infiltration of inflammatory cells, heparin-binding epidermal growth factor (*HBEGF*), a molecule thought to be important in alveolar wall injury/repairs situations [44], CCAAT enhancer binding protein (*CEBPD*), a transcription factor involved in cellular differentiation, proliferation, and inflammation [45, 46], *PIM1*, a gene encoding a serine/threonine kinase with a role in cell survival and proliferation [24, 47], and finally a groups of growth arrest and DNA damage transcripts, *GADD45 α* and *β. GADD45α* when overexpressed in HeLa cell lines caused a significant G_2_-M arrest. Furthermore, overexpression of this molecule causes a reduction of nuclear cyclin B1 protein and subsequent inhibition of cdc2/cyclin B1 kinase activity. Both α and β forms of GADD45 can induce p38/JNK activation and apoptosis [48].

### Transcriptional responses to *P. carinii* life forms are both contact and non-contact dependent

As noted in Table I., a total of 24 genes were expressed greater than 4-fold over the control in *P. carinii* organisms separated from direct contact with A549 cell lines using 0.45 μM pore Transwell inserts. This infection condition was created to provide insights as to which transcriptional responses are created by specific organism to host epithelial cell contact as opposed to possible soluble factors released by the viable organism into the media once outside the rat host. Indeed, others have shown that components such as soluble glucans can be released from fungi and cause biological effects on host cells [49, 50]. Since we are not able to achieve long term growth of *P. carinii* under sterile culture, stringent efforts were made to achieve a germ-free environment. We further verified that the transcriptional responses we report are due to *P. carinii* specific soluble components or contact with the organism. In Supplementary Figure 1, Northern blotting shows the monocyte chemotactic protein-1 (*MCP-1*) transcriptional responses to the various *P. carinii* life form contact conditions. *MCP-1* gene expression has been shown previously to be turned on in response to lipopolysaccharide (LPS) [51, 52]. Although the gene response was different from the microarray fold change reported, the same pattern was noted in Northern blotting by *P. carinii* with (Supplementary Figure 1A,) or without (Supplementary Figure 1B.) pre-treatment for 1.0 h with polymyxin B indicating that LPS contamination was not responsible for our transcription profiles reported. Although Transwell inserts prevent the larger *P. carinii* organisms from contacting the A549 cell lines, the 0.45 μM pore size filter does not exclude possible LPS contamination of the epithelial cell line, hence we believe this represents an important control experiment for our observations.

### GADD34 induction to *P. carinii* trophic forms is time course dependent

We have shown above that the known apoptotic regulator *GADD34* is up-regulated greatest in isolated trophic forms in contact of A549 cells by approximately 52.0-fold versus the control. Overexpression of GADD34, a member of a growing family of growth arrest and DNA damage (hence the designation GADD) proteins previously found to be inducible by various DNA damaging agents [53], has been linked to cell cycle arrest, via activation of p21/WAF1 and accumulation of p53 in the nucleus. In human cells *GADD34* mRNA expression is induced by ionizing irradiation and is maximally expressed at 8 hours but decreases to pre-stimulated levels by 12 hours GADD34 [54].

To further analyze one aspect of gene responses discovered in the microarray experiments related to cell cycle arrest upon *P. carinii* contact with A549 cells, we isolated separate populations of trophic forms (the life form that gave us the greatest fold change induction of *GADD34*) and applied them to A549 cells as stated for the microarray experiments. A time course of *GADD34* expression was determined by Northern blot. Similar to previously published data for induction of *GADD34* by DNA damaging agents such as irradiation in the human ML-1 cell line, *GADD34* expression increased over time, peaking in the 3-to-6-hour range, and decreasing to near basal levels by 12 hours (Figure 3).

**Figure 3.**
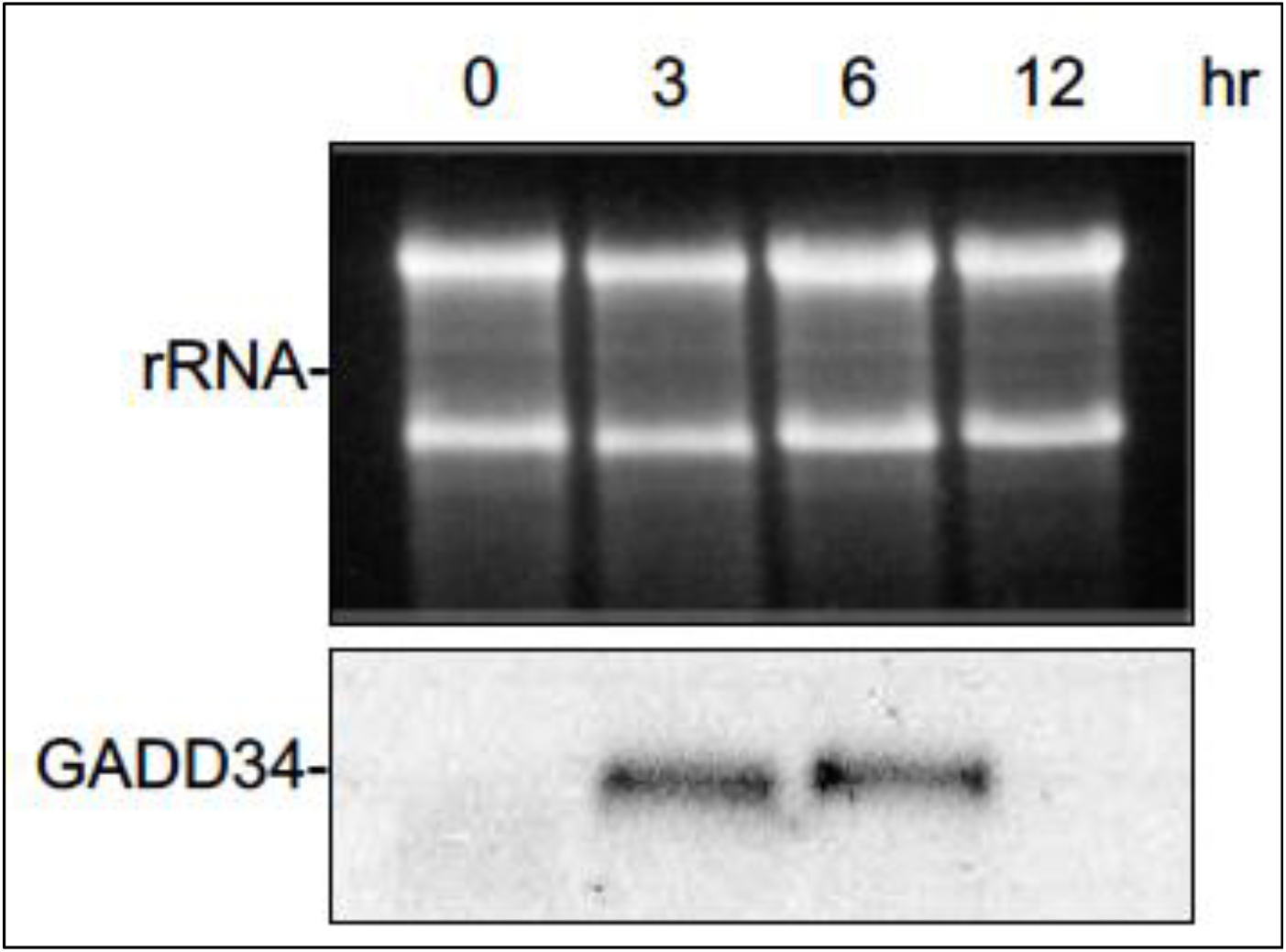
Time course of *GADD34* induction by *P. carinii* trophic forms on A549 cells. Isolated trophic forms of *P. carinii* were applied to A549 cells and total RNA was harvested at the indicated time and analyzed by Northern blotting. The top photograph demonstrates the two A459 cell major ribosomal subunits demonstrating equal RNA loading.

### *P. carinii* contact with A549 cells causes NF-κB activation of the pro-survival/anti-apoptotic BFL-1/A1 promoter

By microarray analysis, we showed that *BFL-1/A1* a member of the pro-survival /anti-apoptotic Bcl-2 family was upregulated upon cyst and trophic form as well as mixed *P. carinii* populations contact with A549 cells (Table II-IV.). Other researchers have shown that *BFL-1/A1* lies downstream of and is a transcriptional target of NF-κB and can block TNFα-induced apoptosis [35]. To determine if the *BFL-1/A1* A549 transcriptional response to *P. carinii* was NF-κB dependent we applied mixed life form *P. carinii* populations to A549 cells transfected with the wildtype *BFL-1/A1* promoter. *P. carinii* contact was allowed to occur for 6 hours (Previously, this time point had shown to give a high-fold gene expression by TNFα-inducible activation of the *BFL-1/A1* promoter.) [35]. Interestingly, we found that the *BFL-1/A1* promoter gave an approximate 3 to 6-fold increase in CAT expression compared to *P. carinii* cultured in Transwell inserts (Figure 4).

**Figure 4.**
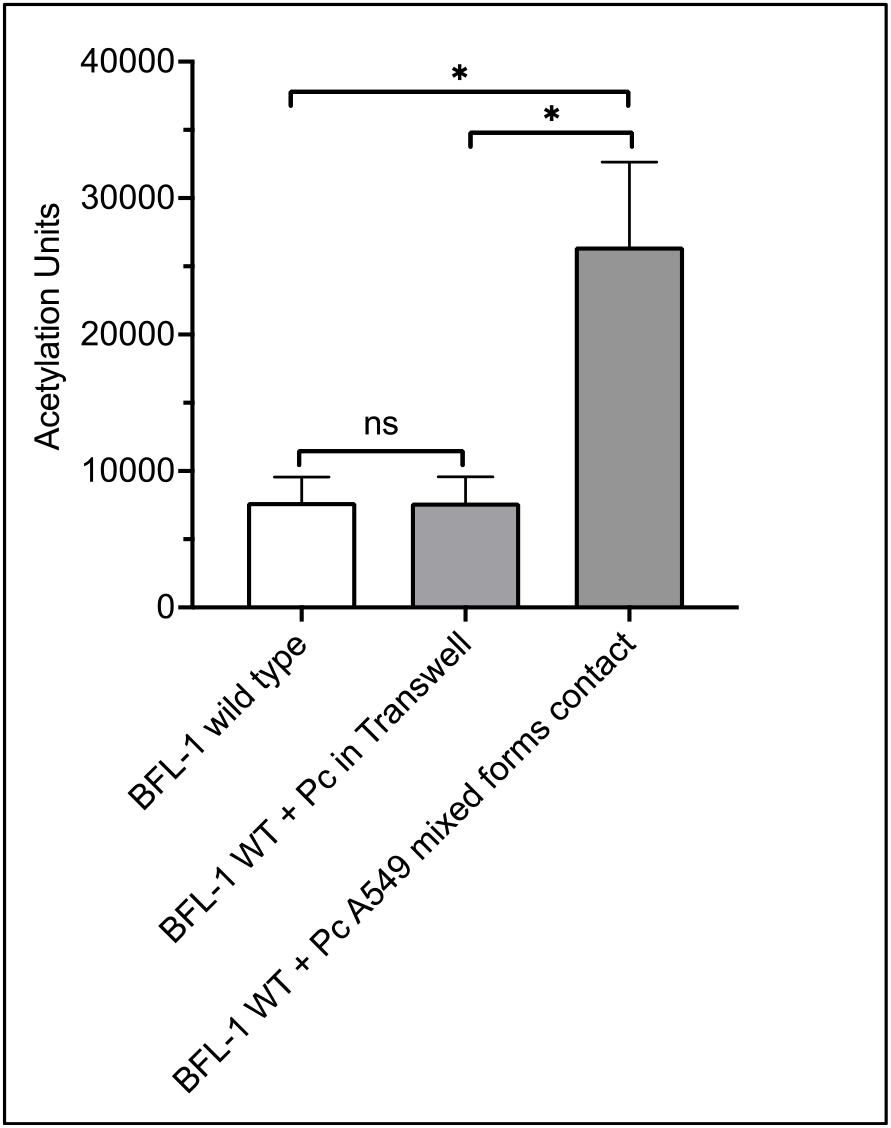
The human *Bfl-1/A1* promoter is induced in A549 upon contact with *P. carinii* (Pc) and is NF-κB dependent. Schematic representation of CAT reporter gene activity as measured by acetylation units. The average acetylation units by CAT from three independent experiments are shown. A549 cells were transfected with wild-type –1374/+81 Bfl-1/A1-CAT [35]. Empty CMV vector was used as a negative control. The bar graph represents the results from three independent experiments. * *P* < 0.05.

Therefore, it appears that *P. carinii* can induce at least one member of the Bcl-2 pro-survival family members in A549 epithelial cells and that the *BFL-1/A1* transcription is controlled by NF-κB proteins.

## CONCLUSIONS

As a continuing step in understanding the complex biological effects of *Pneumocystis* infection, we used high-density microarrays to survey genome-wide transcriptional responses in A549 epithelial cells upon contact with isolated cyst and trophic forms of *P. carinii* as well as mixed populations of the organism. In this study, the response of A549 cells to specific life forms as well as mixed organisms was characterized and found that within 3 hours, infected cells rapidly responded to infection by upregulating the expression of a number of genes. Overall, this response by A549 cells to *P. carinii* infection was dominated by the upregulation of a number of genes associated with the immune response. In addition, we have documented a number of genes upregulated that are life form as well as mixed population specific that occurred following contact with the A549 cell.

As with any model system, our experimental design and reported data have limitations. First, our experiments are limited by data from a single time point as well as a single multiplicity of infection (MOI). We and others have shown that the *P. carinii* MOI of epithelial cell lines can have greatly variable biological responses. For example, applying less than 50:1 *P. carinii* organisms to a mink lung epithelial cell line results in no appreciable effect on p34^*cdc2*^. However, when those ratios of infection exceed 50:1 (e.g., 75-100:1) a significant decrease in kinase activity is observed [55]. Furthermore, when total *P. carinii* organisms from 1 × 10^3^-1 × 10^8^ were applied to rat lung epithelial cells for 4 hours, only the 1 × 10^7^ infection scheme gave a decrease in p38 MAPK signaling [56]. Our system is also limited by one cell type, whereas as in the lung itself, lung epithelial cells modulation would be affected by several other immune and nonimmune cell types as well as soluble factors. Overall, these limitations were accepted to obtain a reproducible, biological model of the interaction of *P. carinii* and a lung epithelial cell line.

To address the specific transcriptional responses of the epithelial cell following contact with *P. carinii*, we included in our experimental design the placement of organisms in a 0.45 μM Transwell insert with equal numbers of mixed *P. carinii* organisms to that of the cell contact condition suspended above the A549 cell monolayer, as well as an untreated control. This addition allowed us to filter out gene responses due to possible soluble factors released from *P. carinii* itself or through its isolation from the immunocompromised rat rather than those gene responses that maybe contact dependent.

Since we cannot grow *P. carinii* under sterile in vitro conditions at the present time, we are unable to be sure that the *P. carinii* obtained from the immunocompromised rat are completely sterile. Very rigorous attempts were made in the experiments to achieve sterile conditions including isolating *P. carinii* from the rat in a laminar flow hood, use of sterile reagents and the discarding of organism preps that appear to have bacteria/fungal contamination. Furthermore, we tested either pretreatment of *P. carinii* organisms with the LPS-binding reagent polymyxin B to determine whether this would alter the observed *MCP-1* gene expression. *MCP-1* transcription has been shown previously to be highly sensitive to LPS [52]. Our results indicate that pretreatment of *P. carinii* organisms with polymyxin B did not change the expression profile, indicating that our sample *P. carinii* preparations appear LPS free.

Analysis of the mRNA expression status of approximately 18,400 human genes demonstrated that *P. carinii* isolated life forms as well as mixed population infections of the A549 cell line induces significant changes in gene expression in a relatively small amount of the completed genes tested (i.e., < 0.5 % according to our cut off of reporting only genes with 4-fold changes). These results indicate that A549 RNA expression to *P. carinii* is not global in nature but rather narrow and specific. Other researchers have shown that in prokaryotic infections of epithelial cell lines, a similar phenomenon is observed [19].

Our studies revealed a number of genes whose mRNA expression was not known to be previously increased upon *P. carinii* interaction with lung epithelial cells. For several of these genes the findings suggest a model of the lung epithelium as a first alert/defense for infection by *P. carinii* that can provide proinflammatory and chemotactic signals to underlying lung cells and tissues. For example, the chemokine receptor CXCR4, a gene we found up-regulated in A549 cells following contact with *P. carinii* mixed populations (FC: +22.8), along with its ligand CXCL12 play an important role in the development of lymphocyte development, trafficking and function [57]. Previously, it has been shown with high density cDNA arrays that CXCR4, also a coreceptor for HIV-1 is induced upon *Mycobacterium tuberculosis* and that this upregulation leads to increased HIV-1 replication in human macrophages [58]. These authors suggest that these changes may be in part why tuberculosis accelerates the course of AIDS [58]. Others have shown previously that PCP can increase the infection titer of HIV-1 in blood and peripheral mononuclear cells (PBMNs) and also that CXCR4 increased in these cells as assayed by immunohistochemistry [59]. Taken together, these results suggest that induction of CXCR4 by *Pneumocystis* can occur in various cell types and that up-regulation of CXCR4 during PCP/HIV-1 may account for the poor prognosis of patients who develop PCP. Another interesting proinflammatory finding from our microarray experiments was the specific fold change of *ICAM-1* (FC: +6.1) in the *P. carinii* cyst form contact with the A549 cell. We have shown previously that mixed populations of *P. carinii* can induce *ICAM-1* expression in A549 cells, further verifying the current and prior result [13]. It was also shown that TNFα, which up regulates ICAM-1 expression, could interact with fungal cell wall β-glucans [13]. In addition, we have shown previously that it is the cyst form as opposed to the trophic form that has the highest levels of β-1,3 glucan synthetase, the enzyme responsible for synthesizing new cell wall β-1,3 glucan [60]. Therefore, our results taken together, indicate that the *P. carinii* cyst form may be the major contributor to increased expression of *ICAM-1* expression following contact with the epithelial cell line. Lastly, we showed that in mixed *P. carinii* population contact with the A549 cell line, endothelin 2 (*EDN2*) (FC: +20.7) is greatly upregulated. Previously, EDN2 has been implicated in macrophage chemoattraction and activation [61]. It maybe that the lung epithelium secretes EDN2 upon *Pneumocystis* infection to recruit macrophages for host defense.

Among the *P. carinii* induced epithelial genes whose products are not expressed on the cell surface or secreted were several cell cycle related transcription factors, cellular kinases, and phosphatases. Of the phosphatases, MAPK-specific phosphatase *MKP-1* (FC: +9.4 trophic form), has been shown to have a role in inhibiting cell cycle progression. Overexpression of this molecule causes growth arrest in the G_0_/G_1_ phase [43]. This molecule has been demonstrated previously to be upregulated by microarray analysis in *Listeria monocytogenes* infection of the human monocyte THP-1 cell line [17]. Interestingly, we have shown previously that *P. carinii* trophic forms can inhibit p34^*cdc2*^ kinase activity and that this data strongly suggest the antiproliferative activity of *Pneumocystis* on lung epithelium maybe due in part to modification of the p34^*cdc2*^ kinase activity [55]. Also, it has been shown by electron microscopy that the *Pneumocystis* trophic form can bind the host cell very tightly [62]. It has been postulated by some that this close interdigitation may block gas exchange between the neighboring capillaries resulting in poor gas exchange and host cell death [63]. Therefore, it appears that the *Pneumocystis* trophic form may utilize multiple methods to cause growth arrest of the epithelial cell. It would be interesting to determine if specific factors (i.e., virulence factors) on the *Pneumocystis* trophic form or the binding of the trophic form to the epithelial cell itself causes cell toxicity. Our lab plans to examine in the future how MKP-1 driven cell cycle arrest is regulated by *Pneumocystis* trophic forms. Finally, our lab detected upregulation of two p53-inducible stress response genes upon mixed *P. carinii* populations in contact with the A549 cell, *GADD45β* (FC: +4.5) and the transcription factor *ATF3* (FC: +26.2). GADD45β and ATF3 have been demonstrated previously to modulate Cdc2 and p21WAF1 activities leading to G_1_ arrest, cell cycle progression inhibition, and apoptosis [64].

Next, NF-κB leads to the transcription of a wide variety of immune responses, including pro-inflammatory cytokines and chemokines [65] which eventually lead to activation of the innate immune response, recruiting inflammatory cells to the initial site of infection [66]. Previously, our lab reported that a β-glucan component of the *P. carinii* cell wall can activate NF-κB in murine RAW 264.7 macrophages [5]. Other researchers reported that this activation maybe mediated through alveolar macrophage (AM) mannose receptors [67]. We report here for the first time that *P. carinii* contact with A549 cells can also stimulate NF-κB transcription and subsequent modulation of downstream targets. For example, after obtaining a significant fold change of the anti-apoptotic BCL-2 homolog *Bfl-1* upon *P. carinii* mixed populations contact with the A549 cell upon microarray, we next determined if NF-κB is a direct transcriptional target of *Bfl-1* by conducting transfection assays with constructs with the known NF-κB binding site in the *Bfl-1* promoter [35]. We were able to show that upon *P. carinii* mixed population binding to the epithelial cell, activation of the wildtype promoter occurred. How NF-κB activation occurs upon *P. carinii* contact with the lung epithelial cell is unknown. Interestingly, we also found in our microarray experiments that lymphotoxin β (*LTβ*) was induced (FC: +16.3) in the mixed *P. carinii* life form population. LTβ is a member of the TNF cytokine family implicated in development of germinal centers during immune responses in the adult [68, 69]. LTβ upon binding to the LTβ receptor (LTβR) recruits the TNFR-associated factor 3 (TRAF3) [70]. This association presumably through interaction of the cytoplasmic regions induces NF-κB activation [70]. Therefore, as expected, it maybe that the NF-κB host response to *Pneumocystis* may vary according to what cell type it encounters.

In summary, our results show that expression of a number of genes was modulated during *P. carinii* infection of A549 cells. Similar as well as unique host gene expression was noted dependent upon what *P. carinii* life form the epithelial cell came in contact with. Gene expression profiling patterns in A549 cells by *P. carinii* infection provide new information for building a comprehensive framework to understand *Pneumocystis* pathogenesis as well as integrate preexisting information about the pathology of *Pneumocystis* infections.

## Supplementary Materials

**“Microarray Expression of Lung Epithelial Cells Upon Interaction with *Pneumocystis carinii* and its Specific Life Forms Yields Insights into Host Gene Responses to Infection”**

**TJ Kottom, AH Limper, et al**.

**Supplementary Table I.**
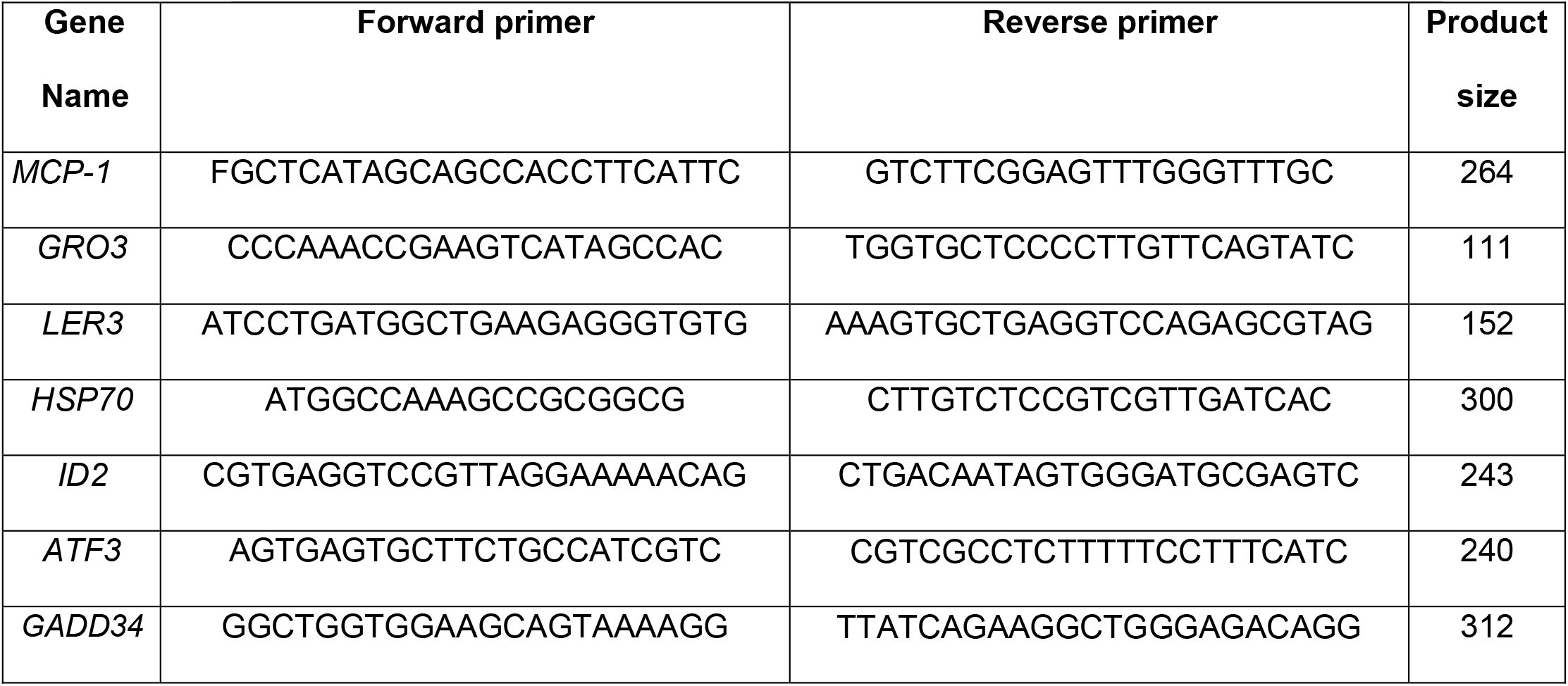
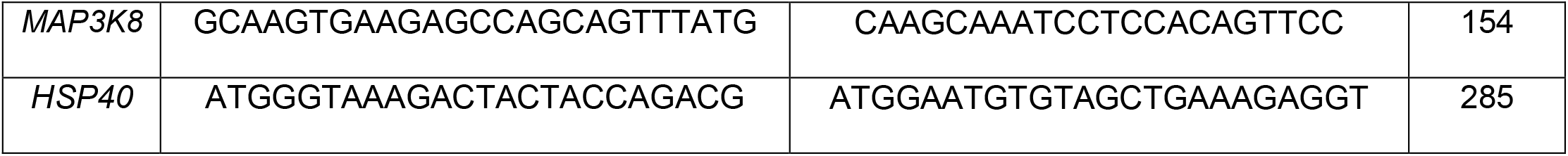
Genes targeted for PCR probe generation and Northern blotting

**Supplementary Figure 1.**
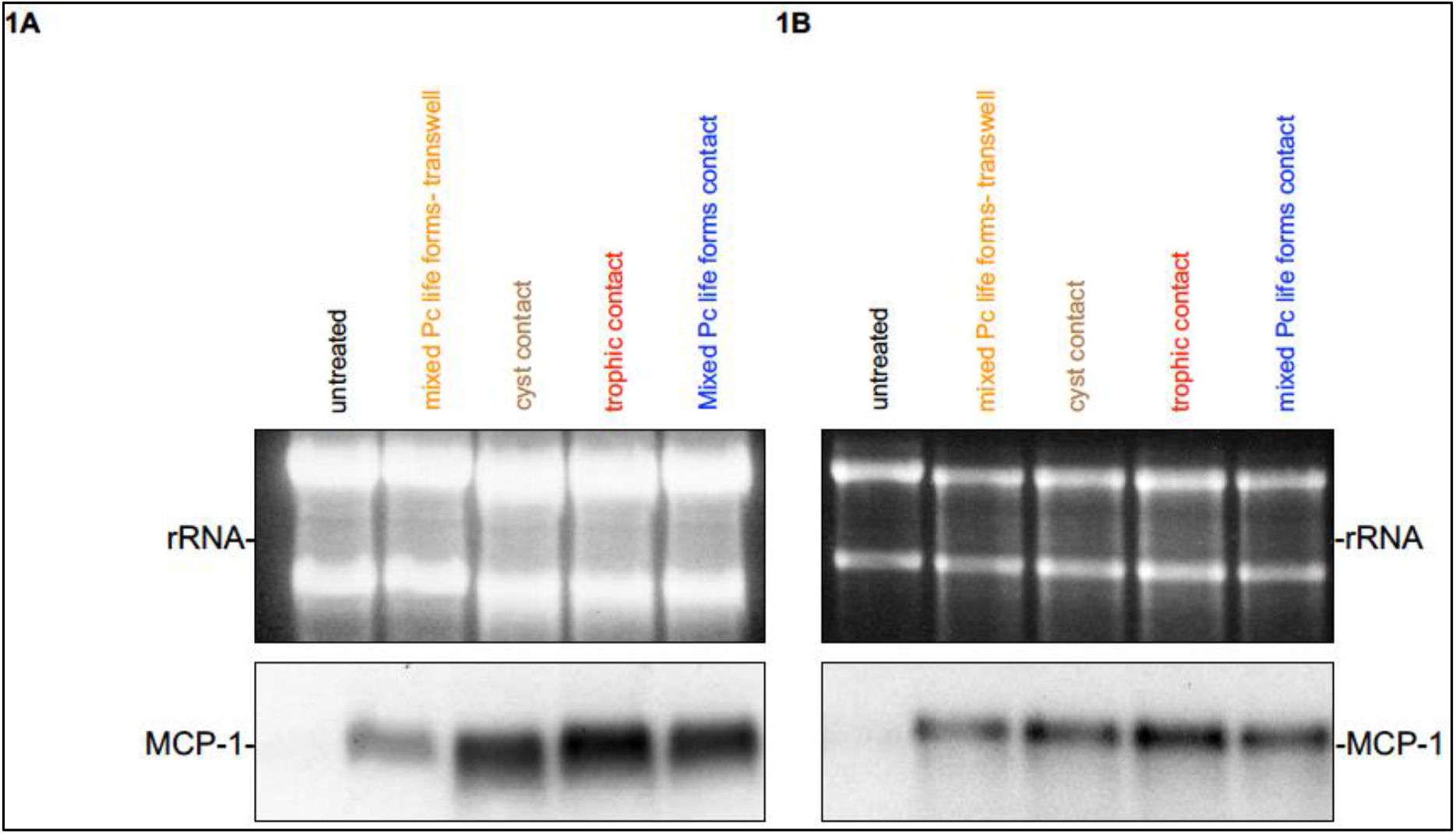
Expression of LPS-inducible monocyte chemotactic protein-1 (*MCP-1*) by A549 cells is similar with (**A**) or without (**B**) pretreatment of *P. carinii* (Pc) with polymyxin B. The top photograph in each panel demonstrates the two A459 cell major ribosomal subunits demonstrating equal RNA loading.

## Notes

**Funding information** This work was supported by the Mayo Foundation, the Walter and Leonore Annenberg Foundation, and NIH grant R01-HL62150 to AHL.

**Conflict of interest** The authors declare that there are no conflicts of interest.

### Competing Interest Statement

The authors have declared no competing interest.

